# *Sarbecovirus*–associated gut microbiome instability in a natural bat reservoir

**DOI:** 10.64898/2026.03.26.714368

**Authors:** Pauline Van Leeuwen, Julia Guillebaud, Marina Voinson, Thavry Hoem, Sreyleak Hoem, Sithun Nuon, Adrien André, Erik Karlsson, Veasna Duong, Julien Cappelle, Johan Michaux

## Abstract

Sarbecoviruses, a subgenus of *Betacoronavirus*, display both respiratory and gastrointestinal tropism, suggesting potential interactions with host gut microbial communities. However, ecological signatures of infection in wild bats remain poorly understood. We investigated associations between *Sarbecovirus* infection status, gut microbiome structure, and diet composition in *Rhinolophus shameli* roosting in northeastern Cambodia. Fecal samples collected across dry and wet seasons (2023–2024) were subject to full-length 16S rRNA gene sequencing and arthropod DNA metabarcoding. *Sarbecovirus–*positive bats exhibited stable alpha diversity but consistent shifts in gut community composition and increased interindividual variability consistent with the Anna Karenina Principle, suggesting infection–associated destabilization of community assembly rather than diversity erosion. Infection status was associated with enrichment of *Shigella* and *Escherichia* species, taxa linked to inflammatory or epithelial stress states in bats. In contrast, dietary composition showed no strong global structuring by infection status and weak coupling with bacterial community structure, suggesting that trophic ecology is unlikely to be the main driver of the infection–associated microbiome signal. Although causal directionality cannot be inferred, our results reveal measurable and consistent microbiome restructuring associated with *Sarbecovirus* detection in a natural reservoir host and highlight the potential of microbiome profiling for monitoring wildlife disease processes.

## Introduction

Coronaviruses are globally distributed RNA-viruses infecting a wide range of species, including humans, and causing a broad spectrum of diseases. Research into the origins of SARS-CoV-2 and continuing interest in *Coronavirus* ecology and evolution have highlighted the value of wild bat surveillance. Bats harbour ancestral lineages of betacoronaviruses from which several viruses of major public health concern have emerged, including SARS-CoV and SARS-CoV-2, both belonging to the subgenus *Sarbecovirus* [1–2]. Despite variability in tropisms of *Sarbecovirus* across bat species [3], intense testing of bats worldwide for betacoronaviruses showed increased detection in rectal or fecal samples compared to other sample types [4]. The key protein responsible for SARS-CoV-2 viral entry into the host cell is glycoprotein S, known as the spike protein. This viral spike protein binds to angiotensin–converting enzyme 2 (ACE2), a cell surface receptor [5] expressed in lung tissue as well as in the oesophageal and intestinal epithelium. As documented in 2020 [6], a variable range of gastro– intestinal (GI) symptoms are observed during SARS-CoV-2 infection in humans. Those symptoms range from diarrhea, nausea, anorexia, abdominal pain and belching [7]. However, the vast majority of studies evaluating the bat host response to *Coronavirus* infection have been performed in cell lines [8], and infected animals did not exhibit evident clinical signs of infection [2, 9]. We could expect that such an infection with GI tract tropism would have consequences on the gut microbiota.

Across wildlife, domestic animals, and humans, reduced microbiome diversity is often associated with pathogenic overgrowth, loss of rare taxa, weakened colonization resistance and increased co-infection risk [10]. Experimental work further suggests that bat-associated gut microbiota may contribute to viral tolerance, as mice transplanted with fecal microbiota from Asian insectivorous bats showed reduced mortality, symptoms, and viral loads following H1N1 infection compared to controls [11].

Host biological traits also shape pathogen dynamics at multiple scales. At the individual level, physiological states such as gestation and lactation involve immunological shifts toward anti-inflammatory responses that may transiently increase viral susceptibility and contribute to sex- and season-specific seroprevalence patterns [12–14]. At the population level, seasonal pulses of viral circulation are largely driven by the synchronous introduction of immunologically naïve juveniles, leading to rapid declines in population) level immunity and increased transmission [12, 15]. Such immune fluctuations may interact with microbial communities, altering community structure and influencing colonization resistance and tolerance mechanisms [16].

Patterns of microbiome variability under infection have led to the application of the Anna Karenina Principle (AKP) to host–associated microbial communities [17]. Under AKP, stressors are expected to induce stochastic, host–specific disruptions of microbial communities rather than a shared, deterministic shift, such that healthy microbiomes are relatively similar to one another, whereas dysbiotic microbiomes diverge among hosts. This manifests as greater β–diversity dispersion rather than a consistent shift in mean composition. In humans with SARS-CoV-2 infection, increased fecal microbiome dispersion correlates with infection status and symptom severity [18, 19]. Evidence in bats remains scarce. A recent investigation in *Artibeus jamaicensis* demonstrated that enteric astrovirus infection altered gut microbial richness in an age-dependent manner yet did not lead to increased among-individual community dispersion [20]. In contrast, infection by *Hibecovirus* in *Hipposideros caffer* reduced bacterial richness and increased interindividual dispersion consistent with AKP predictions [21]. Although causality cannot be confirmed, the short gut transit time of bats and the detectability of viral RNA via qPCR support a potential link between active infection and microbial perturbation [21, 22].

Experimental induction of inflammation in *Rousettus aegyptiacus* also generated AKP-like dispersion and revealed inflammation–associated microbial markers, particularly *Escherichia* spp., consistent with immune–driven alterations of the gut environment [23]. Identifying similar microbial indicators in free– ranging bats may provide valuable, non–invasive insights into subclinical infection or immune activation [9). Beyond pathogens, environmental and ecological factors also influence bat gut microbiomes. Dietary shifts [24–26], habitat disturbance [27–28] and reproductive state [29] can all alter microbial diversity and stability, with potential implications for pathogen dynamics [11). While many bat diets remain to be fully described [30], few studies to date investigated viral shedding with diet composition. Falvo et al. [31] and Vanalli et al. [32] tested experimentally the impact of diet change on influenza viral charge in the Jamaican fruit bat and observed less viral shedding under optimal diet despite not reaching statistical significance in every case explored.

Overall, the literature supports a tight interconnection between bacterial diversity, viral prevalence, host biology, and environmental context. Both causal directions remain plausible: viral infection and associated immune responses may disrupt the microbiome, but pre–existing microbial configurations shaped by diet and seasonality may also influence susceptibility, tolerance, or shedding. Building on this framework, we investigated associations between *Sarbecovirus* infection status, gut microbiome structure and dietary composition in *Rhinolophus shameli*. This Southeast Asian bat species occupies a wide range of forest habitats [33–34] and shows high prevalence of Sarbecoviruses in northern Cambodia [35–36] Yet its ecology is poorly documented, with diet data limited to microscopic analyses indicating predominance of Lepidoptera and Coleoptera [37].

Given the GI tropism of Sarbecoviruses, we hypothesized that *Sarbecovirus* infection would be associated with increased gut microbiome instability consistent with AKP expectations while uninfected individuals would exhibit higher microbial diversity and more constrained community structure. We further hypothesized that diet would act as a major driver of the gut microbiome structure and potentially modulate infection–associated patterns. By comparing bacterial diversity, community composition, and bacterial–diet co-occurrences between *Sarbecovirus*–positive and –negative individuals, we aimed to (i) assess whether infection status is associated with gut microbial imbalance, (ii) evaluate the extent to which such patterns are independent of trophic ecology, and (iii) identify microbial taxa consistently associated with inflammatory or infection-related states in wild bats, without presuming causal directionality.

## Materials and Methods

### 1. Bat capture and sampling

Bat sampling was conducted in Stung Treng province (northern Cambodia) between March 2023 and December 2024 as part of a longitudinal investigation following repeated detections of SARS–CoV–2– related viruses in *Rhinolophus* bats [36, 38]. Nine field sessions were scheduled according to the reproductive phenology of *Rhinolophus* spp., when *Coronavirus* (CoVs) circulation is typically elevated [12]. Sessions occurred every 6–8 weeks from March to August, with an additional late–season survey in 2024. Three previously surveyed hills (Phnom Chhgnauk, Phnom Kar Ngoark and Phnom Chab Pleurng, Figure 1) were repeatedly sampled [36, 38]. Each session lasted six nights, with each site visited twice on non–consecutive nights. Mist nets and harp traps were set at cave entrances and adjacent forested areas.

**Figure 1.**
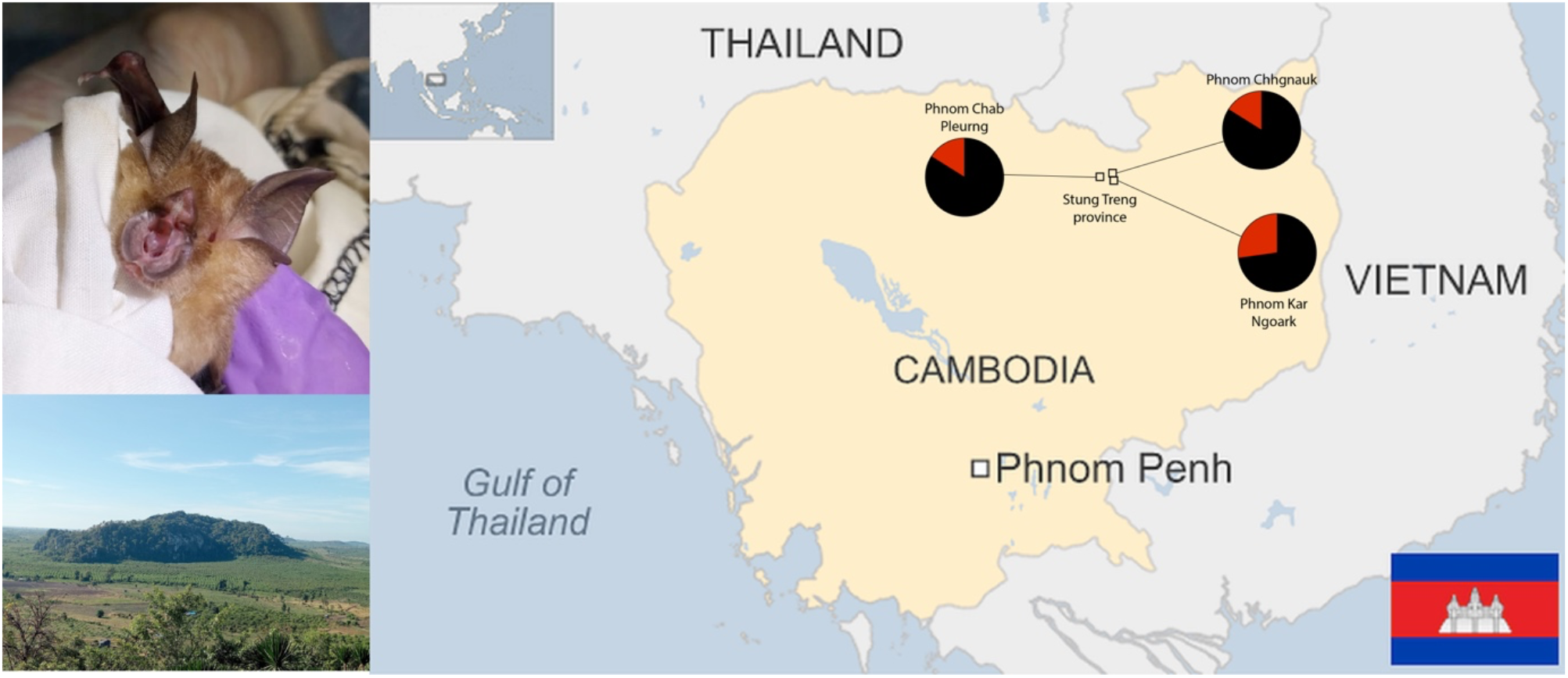
Upper left: *R. shameli* individual (Photo credit: Julien Cappelle), Bottom left: View of Phnom Kar Ngoark (Photo credit: Julia Guillebaud). Right: Map of Cambodia (Source: Wikimedia commons) with sampling sites. Pie charts represent viral status proportion variation between sites (red = *Sarbeco*–positive, black = negative).

Captured bats were individually placed in cotton bag and transported to a mobile field laboratory for species identification and morphometric measurements [39]. Standardized samples were collected for each animal, including rectal swabs (one in viral transport medium, one in TRIzol) and feces. Fecal pellets were preserved in 70% Ethanol and stored on ice until transfer to the Institut Pasteur du Cambodge (IPC). All other samples were kept on ice, transferred to liquid nitrogen within 24 h, and stored at −80 °C at the IPC.

### 2. *Sarbecovirus* screening

*Sarbecovirus* detection followed the protocol described in Guillebaud et al. [36] Total RNA was extracted from TRIzol–preserved rectal swabs using the Direct-zol RNA MiniPrep kit (Zymo Research, USA). A duplex one-step real-time PCR targeting the E and N genes was performed using the Superscript III one–step RT–PCR system with Platinum Taq Polymerase (Invitrogen, Darmstadt, Germany). Samples with Ct < 40 were subjected to Sanger sequencing (Macrogen, Inc., Seoul, Republic of Korea) in both forward and reverse directions using pan-CoV nested-PCR [40] targeting the RdRp gene. The sequences obtained were verified by similarity using the NCBI BLASTn search. A total of 43 bats were *Sarbeco*–positive, 12 CoV–positive and 174 negative individuals (Table 1, Figure 1), and *Sarbecovirus* detected belong to the lineage “group 1,” with representative strains such as RshSTT039 and RshSTT671, related to the earlier RshSTT182/200 viruses [36].

**Table 1.** Sampling list of *R. shameli* individuals according to *Sarbecovirus* qPCR testing, sampling site and session.

| Viral infection status by Site | Sampling session |  |  |  |  |  |  |  |  |  |
| --- | --- | --- | --- | --- | --- | --- | --- | --- | --- | --- |
|  | 03.2023 | 05.2023 | 06.2023 | 12.2023 | 03.2024 | 05.2024 | 06.2024 | 08.2024 | 11.2024 | Total |
| Negative | 22 | 22 | 22 | 21 | 11 | 18 | 21 | 22 | 15 | 174 |
| Chab Pleurng | 8 | 7 | 9 | 12 | 3 | 13 | 7 | 14 | 10 | 83 |
| Chhngauk | 12 | 8 | 4 | 4 | 6 | 2 | 6 | 7 | 2 | 51 |
| Kar Ngaork | 2 | 7 | 9 | 5 | 2 | 3 | 8 | 1 | 3 | 40 |
| *Sarbecovirus* positive |  | 20 | 16 |  |  | 7 |  |  |  | 43 |
| Chab Pleurng |  | 10 | 4 |  |  | 2 |  |  |  | 16 |
| Chhngauk |  | 8 | 9 |  |  | 2 |  |  |  | 19 |
| Kar Ngaork |  | 2 | 3 |  |  | 3 |  |  |  | 8 |
| Total | 22 | 42 | 38 | 21 | 11 | 25 | 21 | 22 | 15 | 217 |

### 3. DNA processing

DNA extraction from feces preserved in ethanol was conducted using the QIAamp Fast DNA Stool Mini Kit from Qiagen following the manufacturer’s protocol. An extraction control was added to each extraction batch (n = 9). Four mock communities from bacterial DNA (HM–782D, BEI Resources) were added to control sequencing accuracy. In PCR_1_, we amplified the full 16SrRNA gene using one primer pair (27F–1492R [41]). Following DNA purification, quantification and pooling at equimolarity, DNA library construction and sequencing were conducted at the University of Liège GIGA Genomics platform using a a total of 217 samples, nine extraction controls, ten PCR_1_ controls and four mock communities samples were sequenced on an GridION (Oxford Nanopore Technologies, Oxford, UK) using a Ligation Sequencing Kit (SQK–LSK112) with 100,000 reads per sample as target.

We used the same DNA extract for the diet of the bats that corresponds to the same samples extracted for microbiome data generation. Briefly, similar to [42] we amplified the mitochondrial cytochrome c oxidase subunit I (COI) using two primer pairs [43–44] Further details on data processing can be found in Supp. material. Following the whole filtering process, our dataset consisted of 1716 distinct biological entities across 153 samples. Due to the low level of taxonomical resolution, we calculated the frequency of occurrence per order for statistical analysis.

### 4. Bioinformatics treatments

After basecalling and barcode demultiplexing using Dorado v1.0.2 and discarding unclassified reads, we controlled the quality of these demultiplexed reads and used *cutadapt* [45] to trim primers for all sequence reads. For the identification of bacteria at species level, fastq files containing full length 16S rRNA gene amplicons were uploaded to the EPI2ME desktop agent 16S workflow (version 2020.2.10, ONT) in which each file was classified using the NCBI 16S rRNA gene blast database through Minimap2 [46] sequence alignment program, yielding closed–reference taxonomic profiles without ASV or OTU inference. Exclusion criteria for nanopore reads were an alignment count accuracy <95%, query cover <90%, quality score (QC) score <10 and read length between 1000 and 2000 bp. The *decontam* package in R was used to detect true contaminants present in extraction and PCR1 controls, resulting in 48 true bacterial contaminants. Mock communities’ samples were checked for accurate taxonomic affiliation (Supp. Figure 1). Control samples were removed, and taxa not belonging to the kingdom Bacteria were also removed leaving a total of 3253 bacterial taxa. Finally, due to great variation in sampling depth between samples with a mean of reads by sample of 74210 (SD = 46880), we rarefied data to 10,000 reads per sample based on rarefaction curves, resulting in a final dataset of 199 samples and 2053 bacterial taxa.

### 5. Statistical analysis

#### a. Diet analysis

We analyzed the structure of the bat dietary assemblages based on the presence–absence of arthropod orders detected in each fecal sample. Non–metric multidimensional scaling (NMDS) was performed using the Jaccard distance to visualize differences in dietary composition among samples. The number of dimensions was selected to minimize stress while retaining ecological interpretability. To identify diet order associated with dietary variation, we fitted diet orders onto the NMDS ordination using the *envfit* function (vegan package). In addition, we computed Spearman rank correlations (with Benjamini–Hochberg procedure) between NMDS axis scores and the relative frequencies of each arthropod order to quantify their direction and strength of association with the ordination gradients. We further tested whether *Sarbecovirus* infection status explained differences in dietary composition using a PERMANOVA (permutations n = 9999) based on the same Jaccard distance matrix. To evaluate temporal trends in dietary composition and potential effects of *Sarbecovirus* infection, we modeled the NMDS axis scores as smooth functions of sampling date. Specifically, Generalized Additive Models (GAMs) were fitted separately for the first two NMDS axes (details in Supp. material).

#### b. Microbiome analysis

Based on collinearity investigations (Supp. material), we retained observed species richness (reflecting taxonomic diversity), Simpson index (reflecting evenness) and Faith’s phylogenetic diversity (phylogenetic breadth) for subsequent analyses of bacterial diversity. For each index, we fitted linear mixed–effects models including age group (Adult, immature and juvenile), sex, site of capture, *Sarbecovirus* infection status, and dietary composition (represented by NMDS1 and NMDS2 scores) as fixed effects, while accounting for season and year as a random intercept to control for temporal dependence among sampling sessions. Models were inspected for residual normality and homoscedasticity, and statistical significance of fixed effects was evaluated using Satterthwaite’s approximation (*lmerTest* package). To evaluate whether *Sarbecovirus* infection was associated with shifts in bacterial community composition (i.e., dysbiosis, AKP), community dissimilarity among samples was quantified using three complementary distance metrics: Bray–Curtis, unweighted UniFrac, and weighted UniFrac. Ordinations were visualized using Principal Coordinates Analysis (PCoA) to illustrate patterns of beta diversity. We tested the effects of host and environmental variables on microbiome composition using a PERMANOVA for each distance metric following the model: Distance ∼Age_Group+Sarbeco_status+Sex+Site. Season_Year was used as a stratification variable in the permutation procedure and permutation tests were performed with 9,999 iterations. We also evaluated multivariate homogeneity of variance using the *betadisper* function.

Following methods detailed in Melville et al. [21), We examined the effect of Sarbecovirus infection status on log–transformed body condition using a linear mixed–effects model with sex and site as fixed effects and season as a random intercept. We additionally used a binomial generalized linear mixed– effects model (logit link) to assess associations between *Sarbecovirus* infection status, microbiome alpha diversity indices, and diet composition (NMDS axes values), including capture periods as a random intercept.

Differences in bacterial taxon abundance between *Sarbecovirus* infection statuses were tested using the ANCOM–BC2 framework on non–rarefied reads. The model was fitted at the species level with infection status as a fixed effect and no random effect. Taxa with prevalence below 10% or total library counts below 500 reads were excluded prior to testing. Significance was assessed at α = 0.05 after false discovery rate (FDR) adjustment of *p*–values.

#### c. HMSC model

We analysed the joint responses of insect taxa and bacterial species using Hierarchical Modelling of Species Communities (HMSC), a Bayesian joint species distribution modelling framework implemented in the *Hmsc* R package [47–48]. We modelled two distinct biological communities measured across the same 139 samples: (1) insect occurrence data (15 taxa; presence/absence), and (2) bacterial abundance data (19 taxa showing differential abundance from ANCOM-BC2 analysis; abundance data). The two response matrices were combined column–wise into a single response matrix. Because the taxa belong to different biological groups and require different statistical error models, we assigned each column a specific response distribution: a *probit* distribution for insect presence/absence data and a *lognormal–Poisson* distribution for bacterial abundance data. We ran MCMC chains with a burn-in period and sampling length sufficient to achieve convergence, evaluated using trace plots, effective sample sizes, and potential scale reduction factors (PSRF). Posterior summaries were obtained for model parameters, including species–specific environmental responses, residual species associations, and latent factor loadings. Model performance was assessed using posterior predictive checks, explained variance, and species–specific predictive power measured via Tjur’s R^2^ (for binary responses) and log-normal pseudo-R^2^ (for abundance data). The residual correlation matrix among species was used to infer potential ecological associations.

## Results

### 1. diet variation and viral status

The overall diet of *R. shameli* was dominated by *Coleoptera* (Frequency Of Occurrence, FOO = 0.451), followed by *Hemiptera* (FOO = 0.379), *Lepidoptera* (FOO = 0.353) and *Diptera* (FOO = 0.333), with lower detection of *Diptera* and *Lepidoptera* in 2023 compared to 2024 (Figure 2A). Despite low level of taxonomical resolution, we detected high FOO for the families of *Cicadellidae* and *Rhyparochromidae* (*Hemiptera*, 0.163 and 0.111), *Scabaridae* (*Coleoptera*, 0.098), *Blattelidae* and *Termitidae* (*Blattodea*, 0.084 and 0.071), *Lecithoceridae, Geometridae* and *Crambidae* (Lepidoptera, 0.065) as well as *Limoniidae* (Diptera, 0.063). Among taxa detected at the species level, the *Anatrachyntis simplex* moth was found in 2.6% of samples which is a common pest of cotton, maize, banana, pomegranate [49].

**Figure 2.**
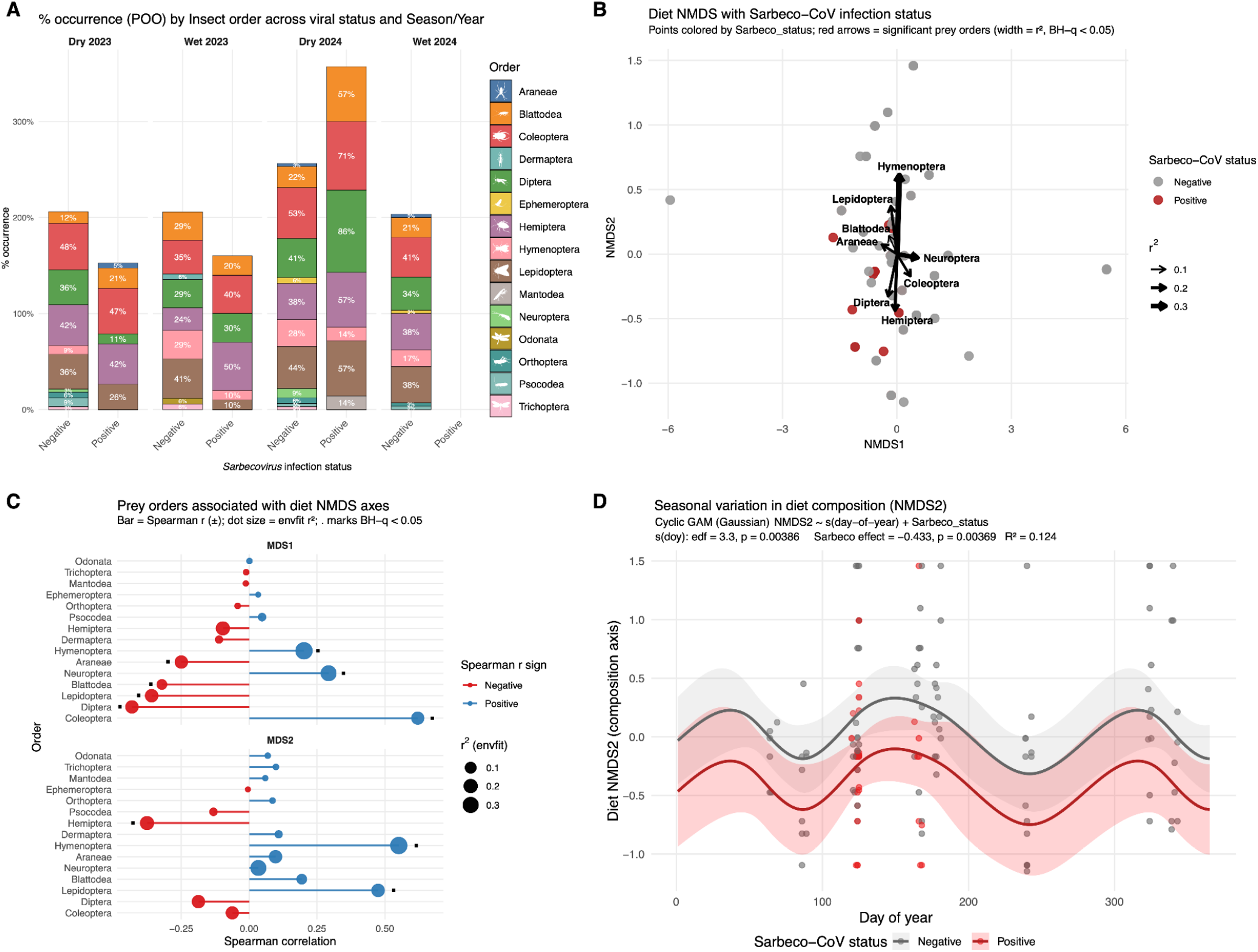
***(A).*** Proportion of occurrence by arthropods order per Season and year in *R. shameli* separated by *Sarbecovirus* infection status detected by PCR. No *Sarbeco*–pos were detected in the wet season of 2024. ***(B)***. NMDS ordination of *R. shameli* dietary composition based on Jaccard distances (stress = 0.095). Each point represents one fecal sample, colored by *Sarbecovirus* infection status. Arrows indicate environmental variables significantly correlated with ordination axes as determined by *envfit*, with arrow length proportional to correlation strength (R^2^ values). ***(C)***. Spearman rank correlations between NMDS axes and relative frequencies of insect orders in *Rhinolophus shameli* diets. The x–axis shows correlation coefficients (*ρ*), and the y–axis lists insect orders. Point color indicates correlation direction (blue = positive; red = negative), and point size is proportional to the *R*_*2*_ of the relationship. Black squares denote significant correlations (*q* < 0.05, Benjamini–Hochberg correction). ***(D)***. Seasonal dynamics of *Rhinolophus shameli* dietary composition modeled by Generalized Additive Models (GAMs). Smooth terms show fitted NMDS2 based on *Sarbecovirus* infection status (red = positive, grey = negative) scores across the day of year (*doy*), with shaded areas representing 95% confidence intervals.

The NMDS ordination based on Jaccard distances (stress = 0.095) provided a robust two–dimensional representation of dietary composition among *R. shameli* individuals (Figure 2B). Samples displayed moderate overlap, suggesting partial differentiation in diet profiles across individuals. The *envfit* analysis confirmed that dietary composition was significantly structured by the relative occurrence of eight insect orders (all *q* < 0.05). The strongest correlates were *Hymenoptera* (R^2^ = 0.37, *p* < 0.001) and *Hemiptera* (R^2^ = 0.19, *p* < 0.001), indicating that variation along the ordination axes primarily reflected contrasts among these dominant prey groups. Correlation strengths were consistent with the *envfit* results (Figure 2C), highlighting a strong positive association between NMDS1 and *Coleoptera* (*ρ* = 0.62, *q* < 0.001) and negative associations with *Diptera, Lepidoptera*, and *Blattodea* (|*ρ*| > 0.30, *q* < 0.01). Along NMDS2, *Hymenoptera* and *Lepidoptera* were positively correlated (*ρ* > 0.45, *q* < 0.001), whereas *Hemiptera* showed a significant negative correlation (*ρ* = –0.38, *q* < 0.001). These patterns confirm that variation in insect order composition primarily reflects contrasts between beetle– and hymenopteran–dominated diets versus those dominated by dipteran or hemipteran prey.

In contrast, *Sarbecovirus* infection status had no detectable effect on overall diet composition (PERMANOVA: R^2^ = 0.01, p = 0.14), and homogeneity of dispersion did not differ between groups (betadisper: p = 0.27). (GAMs were used to assess whether diet composition varied seasonally or in relation to *Sarbecovirus* infection. For the first diet NMDS axis, neither the cyclic smoother for day of year (*edf* = 0.68, *F* = 0.12, *p* = 0.277) nor *Sarbecovirus* infection status (*t* = -0.57, *p* = 0.567) were significant predictors. The model explained only 1.4% of the deviance (adjusted R^2^ = 0.014). In contrast, the second diet NMDS axis displayed a significant seasonal pattern (*edf* = 3.30, *F* = 2.26, *p* = 0.0038) and was negatively associated with *Sarbecovirus* infection (Estimate = -0.43 ± 0.16, *t* = -2.92, *p* = 0.0036). The model explained 14.9% of deviance (adjusted R^2^ = 0.124), suggesting that both temporal variation and infection status moderately contributed to shaping dietary composition along this axis. Specifically, infected bats tended to occupy lower NMDS2 scores, corresponding to a dietary shift towards *Hemiptera* prey, relative to non–infected individuals (Figure 2D).

### 2. Alpha–, beta– diversity in microbiome and viral status

The gut microbial community was dominated by Pseudomonadota (78%; formerly known as *Proteobacteria*) represented mainly by members of bacterial class Gammaproteobacteria (*Enterobacteriaceae, Pasteurellaceae*), and Bacilliota (18.9%; formerly known as *Firmicutes*), by and large ascribed to Bacilli (*Streptococcaceae, Listeriaceae, Bacilliaceae*, Figure 3). Non–infected and Sarbeco–positive bats shared only 26% of bacterial species in common (543 species), representing 92% of the overall microbiome in read abundance (Supp Figure 2).

**Figure 3.**
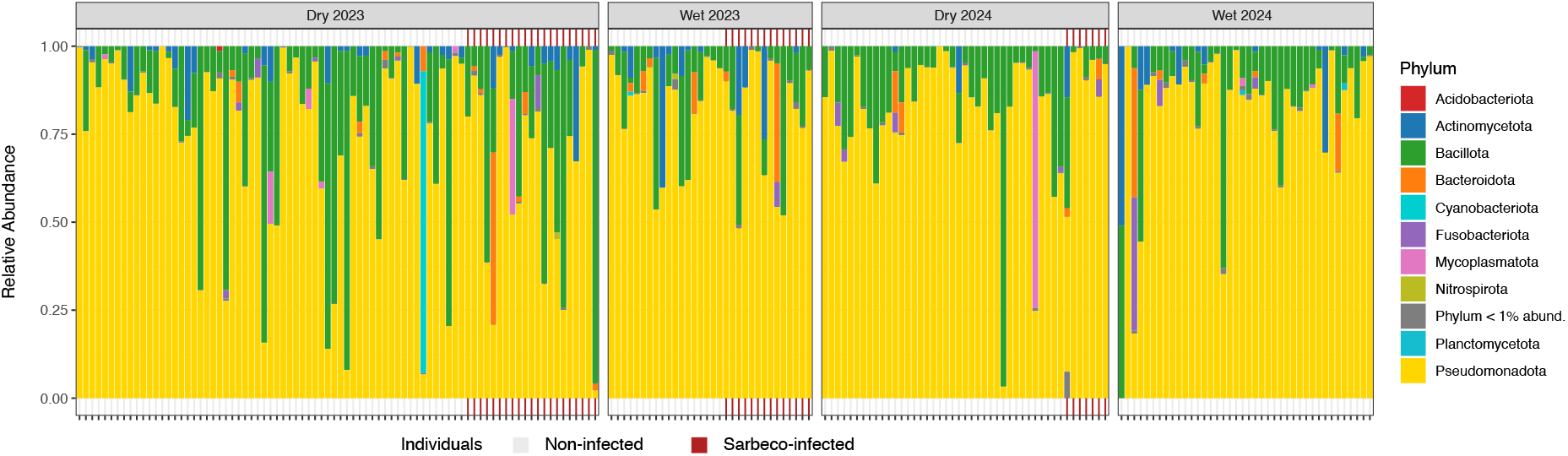
Relative abundance of most common bacterial phyla per *R. shameli* individual across seasonality, with identification of *Sarbecovirus* infection status based on PCR detection.

Linear mixed–effects models were used to explore ecological and host predictors of bacterial alpha diversity. Across all models, the random effect of seasonality explained little variance (SD = 0.04– 12.3), indicating limited temporal influence on bacterial diversity. For species richness, none of the predictors were statistically significant (*p* > 0.05, Supp. Table 1). For Simpson diversity, age group had a modest effect (*p* = 0.002), with immature bats exhibiting lower bacterial evenness (Estimate = -0.09 ± 0.03) compared to adults. No other covariates, including infection status or diet composition, significantly explained variation in Simpson diversity (*p* > 0.24 for all). For Faith’s phylogenetic diversity (PD), the same pattern was found as Simpson index for immature bats (Estimate = -1.29± 0.3, *p* = 0.02), while none of the other tested variables were significant (*p* > 0.10). Overall, bacterial alpha diversity was relatively stable across sex, site, and infection status, with only slight evidence for lower diversity in younger individuals.

Multivariate analyses revealed that *Sarbecovirus* infection was significantly linked to distinct bacterial community composition in bats (Supp. Table 2). Infection status explained a small but significant portion of variation under Bray–Curtis (*F* = 1.97, *R*^*2*^ = 0.01, *p* = 0.006; Figure 4A) and unweighted UniFrac distances (*F* = 1.59, *R*^*2*^ = 0.008, *p* = 0.016), indicating detectable infection–associated dysbiosis. Dispersion tests showed higher within–group variability for infected individuals under Bray– Curtis *F* = 3.75, *p* = 0.05) and weighted UniFrac (*F* = 4.81, *p* = 0.029), suggesting concurrent community instability (AKP, Figure 4B). No consistent effects of age, sex, or site were detected across models (Supp. Table 1). The ANCOM–BC2 analysis identified 19 bacterial species whose relative abundances differed significantly between Sarbeco–positive and Sarbeco–negative *R. shameli* (FDR < 0.05; Figure 5A). Among the most discriminant taxa, *Shigella dysenteriae, S. flexneri, Escherichia coli, E. fergusonii*, and *E. marmotae* were significantly enriched in infected individuals (log–fold change > 1). In contrast, a broad suite of *Enterobacteriaceae* symbionts as well as *Enterococcus casseliflavus* and *Serratia ureilytica* were significantly depleted in infected bats (Figure 5B). Together, these results indicate that *Sarbecovirus* infection in *R. shameli* is associated less with a reduction in within–host bacterial diversity than with a destabilization and reorganization of between–host community structure.

**Figure 4.**
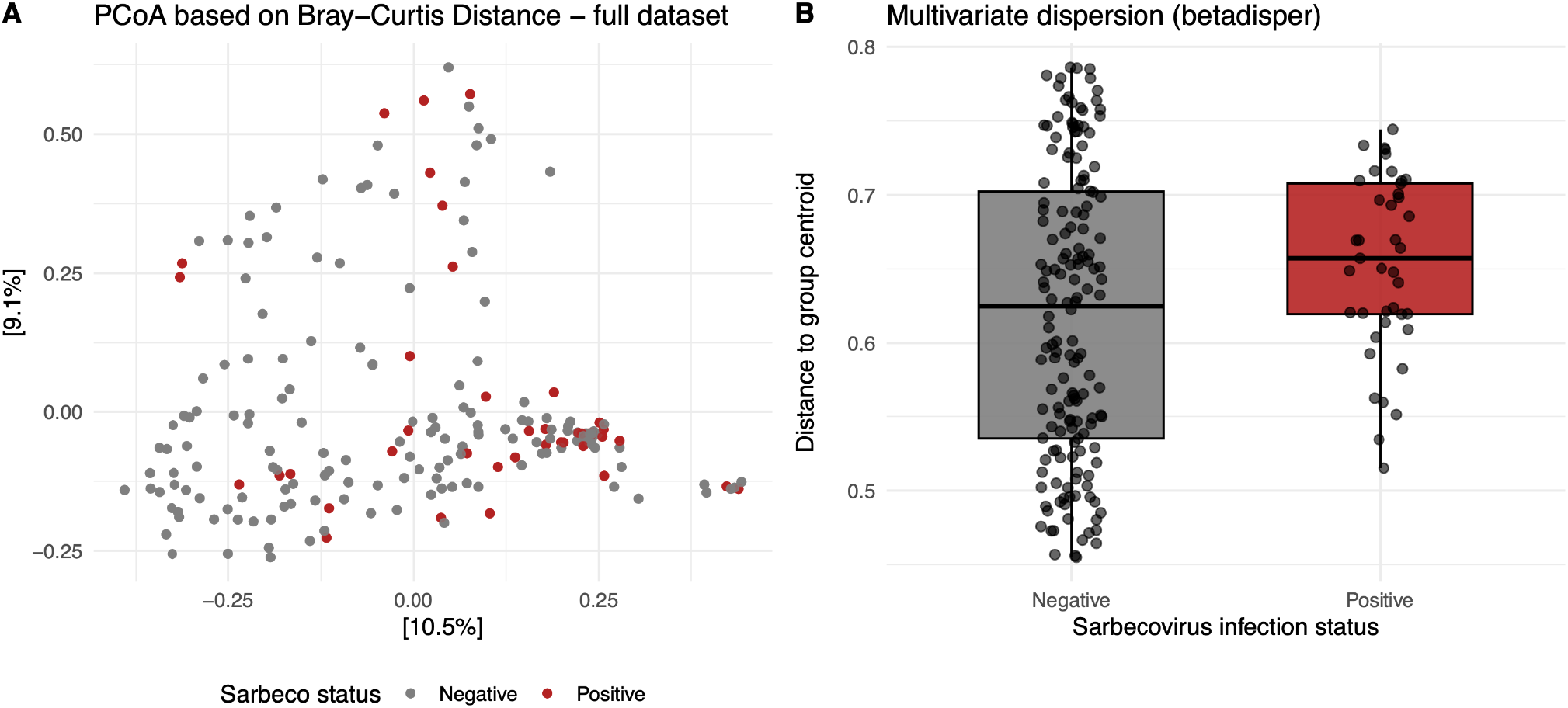
***(A)*** Principal Coordinates Analysis (PCoA) based on Bray–Curtis dissimilarities showing overall community separation between infected (red) and non-infected (grey) individuals. ***(B)***. Multivariate dispersion (betadisper) showing Bray– Curtis distances of samples to group centroids (permutest, *p* = 0.05).

**Figure 5.**
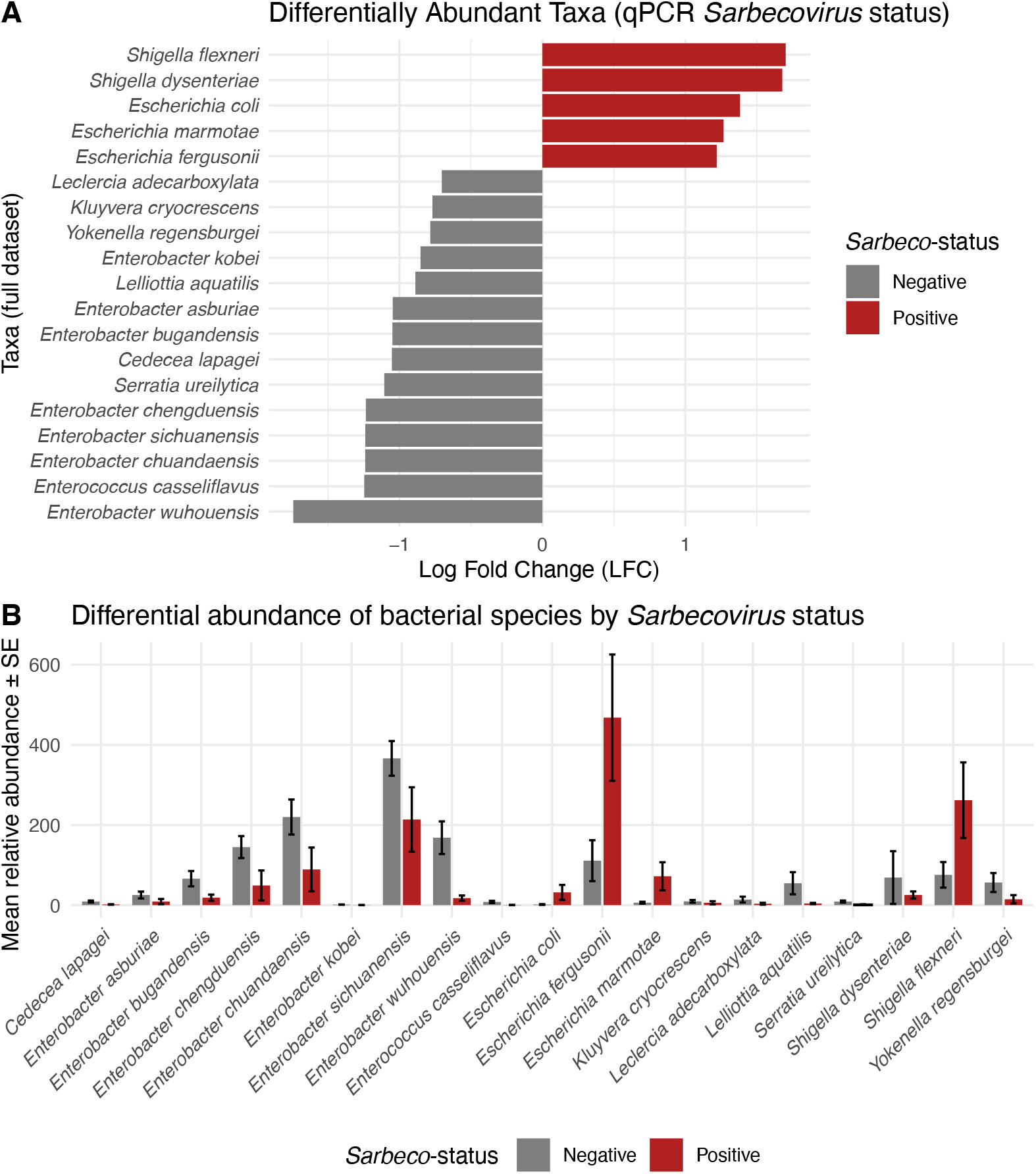
***(A).*** Bias-corrected log fold changes (LFC) from ANCOM–BC2 at the species level comparing *Sarbecovirus*– positive vs. negative bats. Bars show LFC per taxon ordered by LFC magnitude. Multiple testing controlled with FDR (α = 0.05); only significant taxa (q < 0.05) after prevalence (≥10%) filtering are shown. ***(B)***. Mean ± standard error of relative abundance of bacterial species differing between *Sarbecovirus*–positive (firebrick) and negative (grey) *R. shameli* individuals.

### 3. Viral status as response

Among non–pregnant adults (N=113), log body condition did not differ significantly by *Sarbecovirus* infection or sex or site (Supp. Table 3). The random effect of seasonality accounted for a small portion of the variance (SD = 0.026), indicating that body condition remained relatively consistent across sampling periods. *Sarbecovirus* infection status was not associated with microbiome alpha diversity metrics but was significantly related to diet composition along NMDS axis 2 (Supp. Table 4), with increasing values associated with lower odds of infection (odds ratio = 0.48, p = 0.049). suggesting that infected bats may occupy distinct dietary niches with slightly lower NMDS2 scores. The random effect of seasonality explained substantial variation in *Sarbecovirus* infection probability (SD = 1.28 on the log–odds scale), indicating strong seasonal heterogeneity independent of diet and microbiome predictors.

According to the HMSC model, the residual association heatmap revealed strong, coherent structure within the bacterial community and weak structure among arthropod prey. Bacterial taxa formed a highly connected block of strong positive residual correlations, visible as the intense red region in the upper–right portion of the heatmap (Figure 6). This indicates that many bacterial species co–occur more often than expected from shared environmental responses alone, suggesting common unmeasured drivers or facilitative interactions. In contrast, arthropod taxa exhibited only weak and diffuse associations, forming a pale, largely unstructured block at the lower–left of the matrix, consistent with limited residual co–occurrence among arthropods. A notable feature of the matrix is the presence of several bacterial species that show little or no positive residual association with the rest of the bacterial community, visible as pale vertical and horizontal bands interrupting the otherwise strongly correlated bacterial cluster (*Shigella flexneri, Escherichia fergusonii, Escherichia coli, Shigella dysenteriae* and *Escherichia marmotae*, Figure 6). These “weakly connected” bacterial taxa correspond precisely to those that, in a separate analysis, were found to reach the highest abundances in infected samples (Figure 5). Together, these results suggest that infection is linked to the proliferation of a few dominant bacterial species whose dynamics are largely independent of the structure of the wider microbiota or diet composition.

**Figure 6.**
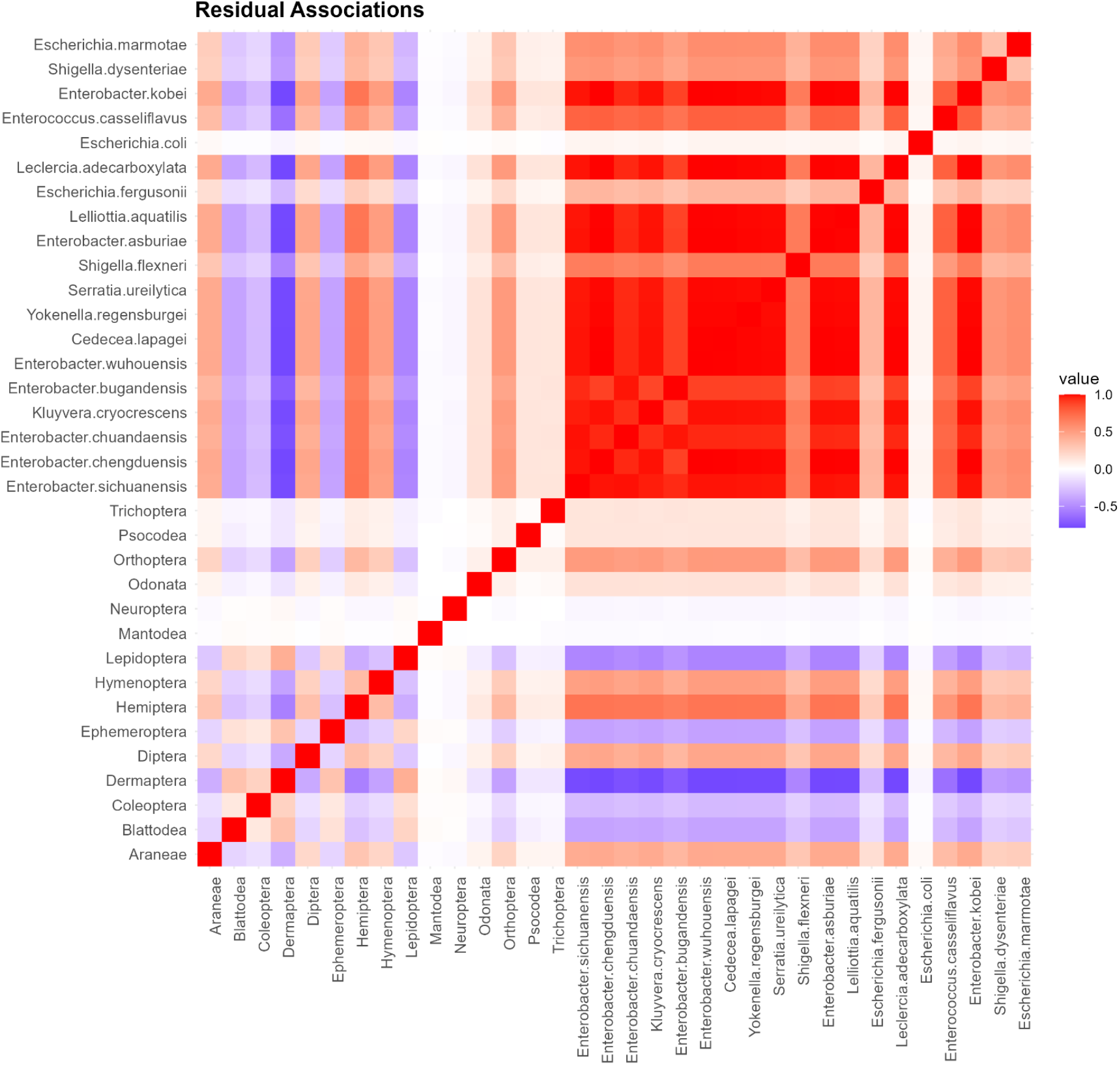
Residual species associations from the HMSC model. The heatmap shows the residual correlation matrix estimated by the HMSC model. Warmer colours (red) indicate positive residual associations, cooler colours (blue) indicate negative associations, and white corresponds to near–zero correlations.

## Discussion

Overall, our study shows that *Sarbecovirus* infection in *R. shameli* is associated with a measurable and consistent but moderate restructuring of the gut microbiome. Most importantly, (i) this signal was characterized by increased interindividual dispersion and compositional shifts rather than by a marked erosion of alpha diversity. In parallel, (ii) diet varied seasonally but showed limited global association with infection status and weak coupling with microbiome structure, suggesting that trophic ecology is unlikely to be the primary explanation for the infection–associated microbial signal. Finally, (iii) infected bats were characterized by enrichment of bacterial taxa previously linked to inflammation or epithelial stress states.

### 1. *Sarbecovirus* infection drives gut dysbiosis in *R. shameli* while alpha diversity remains stable

The predominance of Pseudomonadota in both infected and non–infected individuals is consistent with previous reports from insectivorous bats [27–28]. Contrary to our expectation that infection would be associated with reduced bacterial diversity, alpha diversity metrics remained largely stable across *Sarbecovirus* infection status, sex, and site, with only modest reductions in immature bats. This pattern does not support a simple loss–of–diversity model of dysbiosis and instead suggests that infection– associated microbial responses in this system occur without major erosion of within–host diversity, as documented in other bats [11, 20].

Despite this alpha–diversity stability, *Sarbecovirus* infection was associated with modest but detectable shifts in gut community composition and, importantly, greater interindividual variability. The absence of alpha–diversity loss alongside increased beta–dispersion suggests infection–associated destabilization of community structure rather than diversity erosion. In other words, infected bats did not converge toward a single altered microbiome state; instead, their microbiomes became more heterogeneous, consistent with Anna Karenina Principle expectations for stressed host–associated communities. Infection was associated with compositional differences under Bray–Curtis and unweighted UniFrac, and with increased dispersion under Bray–Curtis and weighted UniFrac, indicating a recurrent infection–associated signal across complementary community metrics. These differences were driven by changes in the relative abundance of dominant taxa, suggesting community-wide reorganization rather than compositional collapse. Similar increases in dispersion were observed following experimentally induced inflammation in *R. aegyptiacus* [23], suggesting that immune or epithelial stress responses may disrupt microbial assembly processes. This interpretation is strengthened by the fact that the signal recurred across complementary beta–diversity approaches, even though infection explained only a small fraction of total variance. In a naturally variable wildlife system, such effect sizes are not unexpected. Field microbiomes integrate multiple sources of variation, including host age, reproductive timing, diet, sociality, microhabitat, and stochastic exposure histories. In that context, a modest but repeated infection signal, coupled with increased dispersion and the enrichment of inflammation–associated taxa, is ecologically meaningful and consistent with a real restructuring of microbial assembly processes rather than statistical noise.

Although causality cannot be established from this cross–sectional design, *Sarbecovirus* detection relied on rectal-swab qPCR and confirmatory PCR/sequencing, which most plausibly reflects active or very recent infection or shedding rather than long–past exposure. This temporal proximity between viral detection and microbiome sampling increases the plausibility that the observed microbial signal reflects infection–associated physiological perturbation, even if reverse or bidirectional relationships remain possible.

### 2. Diet composition does not predict microbiome structure

The diet of *R. shameli* varied seasonally and was dominated by *Coleoptera, Hemiptera, Lepidoptera*, and *Diptera*, in line with previous descriptions of insectivorous *Rhinolophus* diets in Cambodia [37]. This trophic profile reflects the bat’s foraging strategy in dry deciduous and mixed evergreen forests of northern Cambodia [33–34], where beetles and true bugs are abundant in the understory during both wet and dry seasons. The diversity of detected prey included representatives of families *Cicadellidae, Rhyparochromidae*, and *Scarabaeidae*, suggesting that *R. shameli* exploits a wide range of arthropod microhabitats, from foliage to soil detritus. However, *Sarbecovirus* infection did not produce a strong global shift in overall diet composition, as shown by the non–significant PERMANOVA and the absence of dispersion differences between infection groups. Although infected bats showed lower values along the second dietary NMDS axis, this effect was modest and occurred against a broader background of seasonal dietary turnover. Infected individuals may still exhibit distinct foraging preferences or altered prey consumption linked to seasonal availability or physiological state [50].

More importantly, diet and microbiome structure were only weakly coupled. Mantel and Procrustes analyses did not support meaningful covariation between dietary and bacterial community matrices, and the HMSC model revealed weak residual structure among arthropod prey compared with the much stronger structure observed among bacterial taxa. Taken together, these results suggest that trophic ecology is unlikely to be the primary driver of the infection–associated microbiome signal. Rather, diet appears to vary seasonally, while the microbiome signal associated with *Sarbecovirus* detection more likely reflects infection–linked or host–physiological processes [22].

### 3. Inflammation–associated bacterial taxa differentiate infection status

Differential abundance testing using ANCOM-BC2 identified 19 bacterial species differing significantly between infection groups. Similar patterns were found in the HMSC model that included the diet data, strengthening the link between bacterial association with *Sarbeco*–status, regardless of diet items. The consistent enrichment of *Shigella dysenteriae, S. flexneri, Escherichia coli, E. fergusonii*, and *E. marmotae* in infected bats aligns with inflammation–linked taxa reported in both human and experimental bat studies [23–24]. In contrast, *Enterobacter* species, *Enterococcus casseliflavus*, and *Serratia ureilytica* were depleted, indicating loss of potential mutualists involved in mucosal homeostasis. This dual pattern of opportunistic enrichment and symbiont depletion points toward an inflammatory shift in the gut ecosystem of infected individuals, and mirrors microbiome alterations reported in human SARS-CoV-2 infections [18, 23], where *Escherichia–Shigella* enrichment was linked to intestinal inflammation and epithelial barrier disruption. The parallel enrichment of these inflammation–associated genera in *R. shameli* suggests that *Sarbecovirus* infection could provoke subclinical gastrointestinal dysbiosis in bats, consistent with the known enteric tropism of Sarbecoviruses [3]. However, such taxa should be interpreted as indicators of host physiological state rather than direct drivers of pathology. Their enrichment may reflect immune–mediated changes following infection rather than causal involvement in viral susceptibility. The absence of strong correlations between infection and body condition or alpha diversity, as well as the absence of visual symptoms during capture, further supports the idea that these microbial shifts occur under subclinical infection and reflect tolerance mechanisms typical of bat hosts [11].

Several limitations of this study should be acknowledged. First, the cross-sectional design precludes inference on causal directionality between infection and microbiome dysbiosis. Second, viral RNA detection by qPCR most likely reflects active or very recent infection/shedding, but it cannot distinguish ongoing replication from very recent post–infection shedding. Third, although age effects were considered, limited sample sizes constrained our ability to fully resolve age-by-infection interactions previously reported in other bat systems. Finally, the lack of functional immune or metabolomic data limits mechanistic interpretation of inflammation–associated microbial shifts.

## Conclusions

While causality cannot be resolved, the combination of qPCR-based viral detection, gastrointestinal viral tropism, stable alpha diversity, AKP–like increases in beta-dispersion, enrichment of inflammation–associated taxa, and weak diet–microbiome coupling supports the interpretation that *Sarbecovirus* detection is associated with destabilization of gut microbial community assembly in *R. shameli* rather than simple diversity loss. An alternative, non–exclusive interpretation is that pre– existing gut microbiome configurations influence host susceptibility or tolerance to *Sarbecovirus* infection by modulating baseline immune responses, thereby affecting viral replication intensity and detection probability. These results suggest that microbiome instability is more likely linked to host physiological responses to infection than to trophic ecology, without implying strict causality. These findings suggest that infection–associated microbiome instability is more closely linked to host physiological responses than to trophic ecology, while still leaving open the possibility that pre-existing microbiome configurations influence susceptibility or tolerance. Integrating such microbiome profiling into wildlife viral surveillance may therefore improve understanding of host–virus coexistence and tolerance mechanisms [51].

## Acknowledgments

We would like to acknowledge authorities from the Ministry of Agriculture, Forestry, and Fisheries for their support in facilitating this work, and particularly the Department of Wildlife and Biodiversity under the Forestry Administration for their support during field data collection, particularly San Sovannary, Eam Sona and Chhun Vanna. We would like to warmly thank for their cooperation the local authorities and the communities of the Thalaborivat district in Stung Treng province, Cambodia. We also acknowledge the help of Cedric Marsboom (AVIA–GIS), Morgane Labadie (CIRAD), as well as Anaïs Bompard from INRAE, Neil Furey, Tey Putita Ou, Vibol Hul and the entire virology unit of IPC.

## Ethics approval

All procedures complied with relevant national and institutional guidelines for the care and use of animals. Handling and sampling were performed by trained personnel in accordance with the Guidelines [52] of the American Society of Mammologists for the use of wild mammals in research and education, and with the statutory authorization of the Forestry Administration (Ministry of Agriculture, Forestry and Fisheries, Cambodia). The Forestry Administration oversaw all field activities, as no animal ethics committee exists in Cambodia. This study is reported in accordance with ARRIVE guidelines [53].

## Data availability

All data supporting the findings of this study are publicly available upon publication. This includes: (i) sample–level metadata; (ii) a frequency-of-occurrence matrix of arthropod orders per sample derived from DNA metabarcoding; (iii) the bacterial species abundance matrix generated from full–length 16S rRNA sequencing; and (iv) the corresponding taxonomy file containing representative sequences and tree. These datasets are deposited in an open-access repository (https://figshare.com/s/6f68f9e937d1988e93d5), and accession links will be provided in the final version of the manuscript. Additional code used for data processing and analysis will be made available upon request.

