## Supplementary material for "*Sarbecovirus*–associated gut microbiome instability in a natural bat reservoir": supp.

**Supplementary materials**

***Supplementary information: Methods***

*Bacterial DNA PCR information*

PCR_1_ reactions were carried out using Q5 HiFi NEB polymerase, in a final volume of 25 μL containing 0.25 μL of 0.02 U/μL Q5 High–Fidelity DNA polymerase, 0.5 μL of 10 mM dNTPs, 1.25 μL of each of the 0.5 μM forward and reverse primers, 5 μL of 5x Q5 buffer, 5 μL of DNA extract and additional molecular grade water. The thermocycler optimized conditions included an initial denaturation at 98°C for 1 min, followed by 30 cycles of denaturation at 98°C for 20 s, hybridization at 61°C for 30 s extension at 72°C for 2 min, and a final extension at 72°C for 2 min. PCR_1_ negative controls were included in every 96–well plate reaction (n = 10). In PCR_2_, we incorporated a sample–specific combination of 60 bp long in–house tags for Ligation Sequencing Kit (SQK–LSK112). PCR_2_ was carried out in a final volume of 25 μL. A purification step was conducted after each PCR using AMPure XP magnetic beads following the manufacturer’s protocol.

*Diet data generation*

For each primer pair, a total of 212 samples, 9 extraction controls and 10 PCR1 controls were sequenced on an Illumina NovaSeq flow cell using 150 bp paired–end chemistry and targeting 100,000 reads per sample. Trimmed sequence data was imported into QIIME 2 (v2021.8.0, Boylen et al., 2019), where paired–end sequences were denoised, dereplicated into ASVs (Amplicon Sequence Variant) and chimeras were removed using the dada2 plugin. A custom BOLD database was curated using the method described by O’Rourke with some modifications (O’Rourke et al., 2020) and imported into the QIIME 2 environment to be trained against each Galan’s and Zeale’s sequences (outputs of dada2) with the fit–classifier–naive–bayes method. Since the Galan primers detect Chordata DNA, it allows for cross–validation of morphological identification.

*Microbiome data analysis*

GAMs were conducted using the *mgcv* package in R. We used day of the year as a cyclic smoother to account for the circular nature of seasonal time and included *Sarbecovirus* infection status as a fixed factor. Models were fitted using restricted maximum likelihood (REML) for smoothness selection. Model diagnostics were assessed using *gam.check()* to verify the adequacy of basis dimension, residual distribution, and model fit.
We calculated multiple alpha diversity indices to describe within–sample diversity of bacteria, including Shannon, Simpson, inverse Simpson, Fisher’s alpha, Faith’s phylogenetic diversity (PD), and species richness. To avoid redundancy among highly correlated metrics, we first assessed collinearity using Spearman correlation matrices, variance inflation factors (VIF), and pairwise scatterplots. Retained indexes between datasets were then assessed for collinearity. Strong pairwise correlations (|ρ| > 0.75) and high VIF values (>10) indicated substantial overlap among diversity indices.

To investigate relationships between diet and microbiome composition, we conducted Mantel tests with Jaccard distance generated from arthropod order distribution, and Bray–Curtis, Unweighted and Weighted Unifrac dissimilarities from the microbiome dataset. No significant associations were detected for any metric, indicating that variation in diet composition did not predict overall microbiome similarity among bats (Bray–Curtis: *r* = -0.010, *p* = 0.59; weighted UniFrac: *r* = –0.003, *p* = 0.52; unweighted UniFrac: *r* = 0.036, *p* = 0.12). Procrustes analysis was used to assess congruence between diet composition (Jaccard NMDS) and gut microbiome structure (Bray–Curtis PCoA). The analysis minimized squared distances between corresponding samples after rotation and scaling, and significance was tested with 9,999 permutations (PROTEST, *vegan* package). The alignment showed a low Procrustes correlation (*r* ≈ 0.16, *m*² = 0.98, RMSE = 0.085) and was not statistically significant (*p* > 0.05), indicating no meaningful covariation between diet and microbiome ordination spaces.

***Supp. Figure 1.*** Barplot of relative abundance of mock communities.

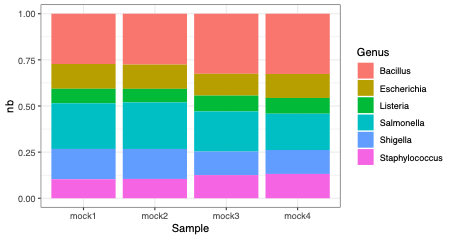

***Supp. Figure 2.*** Euler plots representing (A, C, D) shared bacterial species species between bats based on (A) Sarbecovirus infection status, (C) season and year sampled and (D) sites. B take into account the abundances of each species rather than absence/presence based on viral infection status.

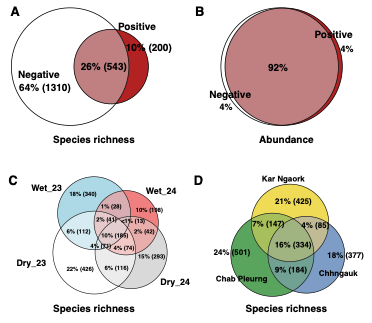

***Supp. Table 1.*** Linear mixed–effects models result of alpha diversity indices.

| **Response variable** | **Predictor** | **Estimate** | **SE** | **df** | **t** | **p–value** | **Random effect** | **Variance** | **SD** |
| --- | --- | --- | --- | --- | --- | --- | --- | --- | --- |
| Species richness | Intercept | 126.361 | 46.310 | 123.668 | 2.729 | 0.007 ** | Season_Year (intercept) | 150.8 | 12.28 |
|  | Age group: Immature | –12.380 | 9.399 | 123.885 | –1.317 | 0.190 |  |  |  |
|  | Age group: Juvenile | 8.546 | 23.504 | 123.792 | 0.364 | 0.717 |  |  |  |
|  | Sex: F | –41.537 | 46.645 | 122.284 | –0.890 | 0.375 |  |  |  |
|  | Sex: M | –39.506 | 46.752 | 122.526 | –0.845 | 0.400 |  |  |  |
|  | Site: Chhngauk | –4.240 | 9.065 | 123.234 | –0.468 | 0.641 | Residual | 1999.0 | 44.71 |
|  | Site: Kar Ngaork | 6.392 | 10.660 | 122.056 | 0.600 | 0.550 |  |  |  |
|  | Sarbecovirus status: Positive | –5.300 | 9.589 | 123.678 | –0.553 | 0.581 |  |  |  |
|  | Diet NMDS1 | –5.658 | 3.595 | 123.617 | –1.574 | 0.118 |  |  |  |
|  | Diet NMDS2 | –3.635 | 6.146 | 123.322 | –0.592 | 0.555 |  |  |  |
| Simpson index | Intercept | 0.88771 | 0.14698 | 123.728 | 6.040 | <0.001 *** | Season_Year (intercept) | 0.001439 | 0.03793 |
|  | Age group: Immature | –0.09105 | 0.02983 | 123.848 | –3.052 | 0.003 ** |  |  |  |
|  | Age group: Juvenile | 0.02358 | 0.07460 | 123.742 | 0.316 | 0.752 |  |  |  |
|  | Sex: F | –0.07204 | 0.14812 | 122.363 | –0.486 | 0.628 |  |  |  |
|  | Sex: M | –0.05617 | 0.14845 | 122.604 | –0.378 | 0.706 |  |  |  |
|  | Site: Chhngauk | –0.03236 | 0.02878 | 123.309 | –1.124 | 0.263 | Residual | 0.020161 | 0.14199 |
|  | Site: Kar Ngaork | –0.02269 | 0.03385 | 122.130 | –0.670 | 0.504 |  |  |  |
|  | Sarbecovirus status: Positive | –0.03628 | 0.03043 | 123.598 | –1.192 | 0.236 |  |  |  |
|  | Diet NMDS1 | 0.01009 | 0.01141 | 123.673 | 0.884 | 0.378 |  |  |  |
|  | Diet NMDS2 | –0.01508 | 0.01951 | 123.386 | –0.773 | 0.441 |  |  |  |
| Faith's PD index | Intercept | 8.29814 | 2.70878 | 123.984 | 3.063 | 0.003 ** | Season_Year (intercept) | 0.1977 | 0.4446 |
|  | Age group: Immature | –1.29035 | 0.54873 | 122.648 | –2.352 | 0.020 * |  |  |  |
|  | Age group: Juvenile | 0.94350 | 1.37148 | 122.271 | 0.688 | 0.493 |  |  |  |
|  | Sex: F | –2.30094 | 2.74509 | 122.936 | –0.838 | 0.404 |  |  |  |
|  | Sex: M | –2.39170 | 2.74996 | 123.227 | –0.870 | 0.386 |  |  |  |
|  | Site: Chhngauk | –0.30038 | 0.53213 | 123.961 | –0.564 | 0.573 | Residual | 6.9463 | 2.6356 |
|  | Site: Kar Ngaork | 0.35437 | 0.62769 | 122.524 | 0.565 | 0.573 |  |  |  |
|  | Sarbecovirus status: Positive | 0.08843 | 0.55882 | 120.500 | 0.158 | 0.875 |  |  |  |
|  | Diet NMDS1 | –0.24926 | 0.21077 | 123.991 | –1.183 | 0.239 |  |  |  |
|  | Diet NMDS2 | –0.33565 | 0.36078 | 123.917 | –0.930 | 0.354 |  |  |  |

***Supp. Table 2.*** PERMANOVA, betadisper, and pairwise comparisons for microbiome composition across datasets.

| **Distance metric** | **Factor** | **PERMANOVA pseudo–F** | **R²** | **p–value** | **betadisper F** | **betadisper p** |
| --- | --- | --- | --- | --- | --- | --- |
| Bray–Curtis | **Sarbeco_status** | **1.969** | **0.010** | **0.0063** | **3.753** | **0.050** |
|  | Age_Group | 1.204 | 0.012 | 0.1800 | 1.649 | 0.195 |
|  | Site | 1.192 | 0.012 | 0.1513 | 0.748 | 0.474 |
|  | Sex | 1.144 | 0.011 | 0.1377 | 0.776 | 0.380 |
| Unweighted UniFrac | **Sarbeco_status** | **1.585** | **0.008** | **0.016** | 0.681 | 0.410 |
|  | **Age_Group** | **1.356** | **0.013** | **0.038** | 1.361 | 0.259 |
|  | **Site** | **1.601** | **0.016** | **0.002** | 0.273 | 0.762 |
|  | Sex | 1.053 | 0.010 | 0.330 | 0.292 | 0.59 |
| Weighted UniFrac | **Sarbeco_status** | **1.178** | **0.050** | **0.292** | **4.805** | **0.029** |
|  | Age_Group | 1.175 | 0.099 | 0.286 | 0.731 | 0.483 |
|  | Site | 1.271 | 0.108 | 0.208 | 0.745 | 0.477 |
|  | Sex | 1.287 | 0.109 | 0.140 | 0.723 | 0.397 |

***Supp. Table 3.*** Effects of *Sarbecovirus* infection status on body condition in adult, non–pregnant bats. Linear mixed–effects model with Season_Year as a random intercept (REML fit). Degrees of freedom and p–values were estimated using Satterthwaite’s approximation.

| **Fixed effect** | **Estimate** | **SE** | **df** | ***t*** | ***p*** |
| --- | --- | --- | --- | --- | --- |
| Intercept | −1.601 | 0.035 | 17.9 | −46.32 | <0.001 *** |
| Sarbeco status (infected vs. negative) | −0.033 | 0.034 | 92.9 | −0.99 | 0.326 |
| Sex (male vs. female) | −0.051 | 0.030 | 99.0 | −1.69 | 0.094 |
| Site: Chhngauk | 0.023 | 0.034 | 96.3 | 0.68 | 0.497 |
| Site: Kar Ngaork | −0.026 | 0.037 | 96.2 | −0.71 | 0.481 |
| **Random effect** | **Variance** | **SD** |  | | |
| Season_Year (intercept) | 0.00072 | 0.027 |  |  |  |
| Residual | 0.02188 | 0.148 |  |  |  |

***Supp. Table 4.*** Diet and microbiome predictors of *Sarbecovirus* infection status. Generalized linear mixed–effects model (binomial error distribution, logit link) fitted by maximum likelihood (Laplace approximation), with Season_Year included as a random intercept.

| **Predictor** | **Estimate (log–odds)** | **SE** | ***z*** | ***p*** |
| --- | --- | --- | --- | --- |
| Intercept | −1.316 | 1.443 | −0.91 | 0.362 |
| Faith’s PD | 0.277 | 0.199 | 1.39 | 0.164 |
| Simpson diversity | −0.585 | 1.667 | −0.35 | 0.726 |
| Species richness (SR) | −0.017 | 0.013 | −1.30 | 0.194 |
| Diet NMDS1 | −0.191 | 0.227 | −0.84 | 0.400 |
| Diet NMDS2 | −0.741 | 0.378 | −1.96 | 0.050* |
| **Random effect** | **Variance** | **SD** |  |  |
| Season_Year (intercept) | 1.627 | 1.275 |  |  |
